# Maternal diabetes induces senescence and neural tube defects sensitive to the senomorphic Rapamycin

**DOI:** 10.1101/2020.09.18.303222

**Authors:** Cheng Xu, Wei-Bin Shen, E. Albert Reece, Hidetoshi Hasuwa, Christopher Harman, Sunjay Kaushal, Peixin Yang

**Affiliations:** Department of Obstetrics, Gynecology & Reproductive Sciences, University of Maryland School of Medicine, Baltimore, Maryland, USA; Department of Biochemistry & Molecular Biology, University of Maryland School of Medicine, Baltimore, Maryland, USA; Department of Surgery, University of Maryland School of Medicine, Baltimore, Maryland, USA; Department of Molecular Biology, Keio University School of Medicine, 35 Shinanomachi Shinjuku-ku, Tokyo 160-8582 JAPAN

**Keywords:** Premature senescence, miR-200c, Forkhead transcription factor 3a, cell cycle inhibitors, maternal diabetes, neural tube defects, Rapamycin

## Abstract

Neural tube defects (NTDs) are the second most common structural birth defects. Senescence, a state of permanent cell cyle arrest, only occurs after neural tube closure. Maternal diabetes-induced NTDs, severe diabetic complications leading to infant mortality or lifelong morbidity, may be linked to premature senescence. Here we report that premature senescence occurs in the mouse neuroepithelium and disrupts neurulation, leading to NTDs in diabetic pregnancy. Premature senescence and NTDs were abolished by deleting the transcription factor *Foxo3a*, the *miR-200c* gene, the cell cycle inhibitors *p21* or *p27*, or by transgenic expression of the dominant-negative FoxO3a mutant or by the senomorphic rapamycin. Double transgenic expression of p21 and p27 mimicked maternal diabetes in inducing premature neuroepithelium senescence and NTDs. These findings integrate transcription- and epigenome-regulated miRNAs and cell cycle regulators in premature neruoepithelium senescence, and provide a mechanistic basis for targeting premature senescence and NTDs using senomorphs.

## Introduction

Neural tube defects (NTDs), the second most common structural birth defects, greatly contribute to infant mortality and lifelong mortality. Folate is the only prevention to NTDs but only prevents up to 70% NTDs ^1^. NTDs are still highly prevalent. Globally, there are 300,000–400,000 cases per year ^2^, resulting in approximately 88,000 deaths and 8.6 million disability-adjusted life years ^3^. Annual medical and surgical costs for children born with NTDs in the US are more than $200 million ^4^. Thus, new therapeutic targets need be revealed for the development of new preventions to NTDs. Majority of human NTDs are caused by non-genetic factors. Maternal diabetes is a major and ever-rising non-genetic factor that induces NTDs ^5^. Approximately three million American and sixty million worldwide women of reproductive age (15-44 years) with pre-gestational diabetes ^6^ make the revealation of the cause of diabetic embryopathy an urgent task.

Using rodent models of diabetic embryopathy, our previous studies have demonstrated that maternal diabetes specifically induces NTDs by altering kinase activation and gene expression ^7, 8,9, 10, 11, 12, 13, 14^. However, how the molecular events downstream of hyperglycemia integrate and lead to failed neurulation is unknown. Neurulation, one of the critical and fundamental morphogenesis processes of embryonic development, leads to the formation of the central nervous system. During neurulation, neuroepithelial cells translocate to the apical side of the neuroepithelium where they undergo chromosome condensation and segregation during mitosis ^15, 16^, and the significance of this feature is unknown. Disrupting this event by senescence in these proferliating neuroepithelial cells may contribute to NTDs.

Cellular senescence in adult cells plays pleiotropic roles in many physiological and pathological processes, including aging, aging-related neurological disorders, tissue repair, tumorigenesis and cardiovascular diseases^17^. Recent studies have revealed that senescence also contributes to embryonic development ^18, 19^. Developmental senescence, a newly discovered, normal programming mechanism, promotes tissue remodeling and plays instructive roles in embryonic development ^18, 19^. Cellular senescence and developmental senescence share a set of features, including senescence-associated β-galactosidase activity (SAβG), heterochromatic foci, and acquisition of a senescence-associated secretory phenotype (SASP) ^18, 19, 20, 21^. However, developmental senescence is unique from cellular senescence because it does not depend on the activation of the cellular DNA damage response or the p16 tumor suppressor pathway, and does not produce some of the SASP factors ^18, 19^. Developmental senescence is mediated by p21 and regulated by the Forkhead transcription factor (FoxO) and TGFβ/Smad pathways ^18, 19^.

Development senescence occurs after neural tube closure at embryonic day 11 (E11) in the mouse ^18, 19^. The cellular senescence inducers, including DNA damage and p53 up-regulation, have been observed in diabetic embryopathy ^22, 23, 24^. Strong evidence also suggests that apoptosis signal-regulating kinase 1 (ASK1), a mediator of cellular senescence induced by high glucose *in vitro* ^25^, plays a causal role in diabetes-induced NTDs ^12^. Finally, maternal diabetes-induced oxidative stress triggers the activation and the increase of cellular senescence mediators ASK1, FoxO3a and p53. Because antioxidants delay or prevent cellular senescence ^26, 27^, oxidative stress may induce senescence. This evidence collectively implies that premature senescence in the developing neuroepithelium may be associated with diabetes-induced NTDs. The developing neuroepithelium is a highly proliferative and dynamic tissue ^28^. Neuroepithelial cells move from basal side to the apical (lumen) side for commencing mitosis ^16, 28^, and, then, translocate in a ventral-dorsal direction in achieving a critical mass for the paired dorsolateral hinge points, which is critical for the two opposite neural folds to be contacted and fused ^28^. Altered neuroepithelial cell proliferation and differentiation, and enhanced apoptosis are contributing factors in NTD pathogenesis ^12, 13, 29^. Thus, it is plausible that premature senescence in proliferative neuroepithelial cells at the apical side would attenuate or collapse the dorsal hinge points, and lead to failed neural fold contact and fusion in diabetic embryopathy.

The present study reveals that maternal diabetes triggers premature developmental senescence (hereafter referred to as premature senescence) in the apical neuroepithelial cells during neurulation, thereby leading to NTD formation. The FoxO3a-miR200c pathway silenced the transcription repressors, ZEB1/2, leading to the up-regulation of the senescence mediators, p21 and p27. Overexpression of p21 and p27 in the developing neuroepithelium mimicked diabetes in inducing apical neuroepithelial cell senescence and NTDs. Since senomorphs and senolytics, which inhibit the ontogeny of senescence and selectively remove senescent cells, respectively, are proposed as a new generation of human disease therapeutics ^30, 31^, we determined the therapeutic effect of the senomorphic rapamycin on diabetes-induced NTDs. Thus, premature senescence in the developing neuroephtium is critically involved in NTD formation in diabetic pregnancy and rapamycin may be an effective therapeutic.

## Results

### Maternal diabetes induces premature senescence in the apical neuroepithelium

We found that mice with maternal diabetes induced a robust staining signal of senescence-associated β-galactosidase (SAβG), a well-recognized senescence marker, in the apical side of the anterior neuroepithelium, on embryonic day 8.5 (E8.5) (Fig. 1A), suggesting that senescence targets a population of proliferative cells in the apical neuroepithelium ^15, 16, 28^. The induction of the major developmental senescence mediator, p21 ^18, 19^, was observed at E8.0 in the same region (Fig. 1B), preceding the SAβG signal at E8.5 onwards (Fig. 1A), suggesting that p21 may trigger senescence in diabetic embryopathy. Cross-sectional analyses through the whole embryo revealed that the SAβG signal was induced by maternal diabetes in the anterior but not the posterior neuroepithelium (Fig. S1), and that the notochord was SAβG positive under both nondiabetic and diabetic conditions (Fig. 1A and Fig. S1). In E9.25, intense SAβG signal was observed in the fusing tips of the two neural folds with a wide gap (Fig. 1C), and contributed to failed neural tube closure in diabetic pregnancy. At an early stage, the dorsal edges of neural folds did not contain senescent cells (Fig. 1A), suggesting that at E9.25, cells at these sites become proliferative and thus susceptible to senescence.

**Figure 1.**
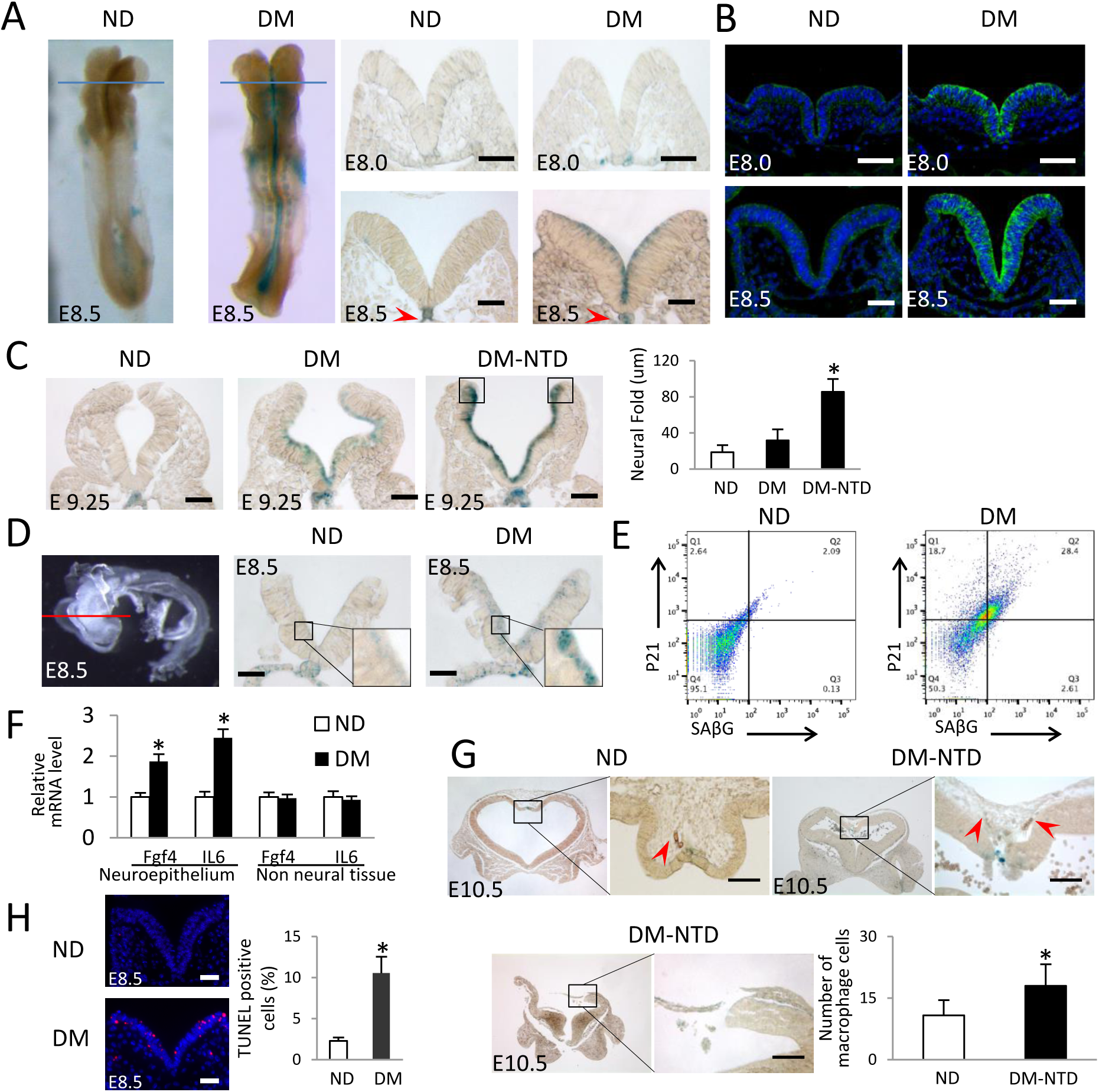
Maternal diabetes induces premature senescence in the developing neuroepithelium. (A) Whole mount staining of senescence associated β-galactosidase (SAβG) in E8.5 embryos and coronal sections of E8.0 and E8.5 embryos after SAβG staining. Blue line indicates the levels of sectioning. Red arrow heads indicated the notochords. (B) Anti-p21 antibody staining on E8.0 and E8.5 embryo sections. (C) Sections of E9.25 embryos after SAβG staining, and quantification of the distance between the two neural fold fusing edges (square frames). (D) Neuroepithelia were isolated from E8.5 embryos by Dispase-II treatment for SAβG staining. Red line indicates section positions. (E) FACS analysis by SAβG and p21 antibody staining in cells of isolated E8.75 neuroepithelia. (F) mRNA abundances of Fgf4 and IL6 in isolated neuroepithelia and non-neural tissues of E8.5 embryos. (G) Sections of E10.5 embryos after SAβG staining. Infiltrated macrophages were indicated by red arrow heads. The graph showed the number of macrophages (n = 3). (H) Representative images of the TUNEL assays in E8.5 embryos and quantification of apoptotic cells. Apoptotic cells were labeled in red. Scale bars = 70 μm in A, B, C, D, H; Scale bars = 45 μm in G. ND: nondiabetic dams; DM: diabetic mellitus dams; NTD: neural tube defects. * indicate significant difference (*P* < 0.05) compared to other groups.

The SAβG signal in the apical side was confirmed in isolated neuroepithelia that avoided the possible X-gal penetration problem during staining (Fig. 1D). Flow cytometry analysis was used to further characterize the features of senescent cells in the neuroepithelium. SAβG positive cells were co-stained with p21 in isolated neuroepithelial cells (Fig. 1E): 92% SAβG^+^ cells were p21 positive, and diabetes increased the number of double stained SAβG^+^ and p21^+^ cells by approximately 14-fold (Fig. 1E). Senescent cells also exhibited the Senescence-Associated Secretory Phenotype (SASP), which adversely impacts neighboring cells. The expression of two SASP factors, interleukin 6 (IL-6) and FGF4, was significantly increased in neuroepithelia of embryos exposed to diabetes (Fig. 1F). Senescent cells are removed by macrophages and induce apoptosis in neighboring cells. Indeed, increased macrophage infiltration in the area of SAβG^+^ cells presence was observed in the defective neuroepithelia of some NTD embryos (Fig. 1G), whereas other NTD embryos completely lost the SAβG^+^ portions of their neuroephithelia (Fig. 1G). Apoptotic cells were mainly present in the SAβG^+^ apical side of the neuroepithelium (Fig. 1H). Thus, maternal diabetes induces premature senescence in the developing neuroepithelium, which resembles the key features of developmental senescence ^18, 19^.

### Deleting the *Foxo3a* gene diminishes premature senescence in diabetic pregnancy

FoxO3a, which is critically involved in developmental senescence ^19^, is activated and participates in maternal diabetes-induced NTDs ^12^. To determine whether FoxO3a is responsible for maternal diabetes-induced premature senescence in the developing neuroepithelium, three types of embryos were examined: Wild-Type (WT), *Foxo3a* deleted, and FoxO3a dominant negative (DN-FoxO3a). *Foxo3a* deletion diminished the signals of SAβG, the DNA damage marker γH2AX and the heterochromatin marker histone H3 trimethylation at lysine-9 (H3K9me3) (Fig. 2A, B). It also restored phospho-Histone H3 (p-H3) signals, which showed that proliferating cells in metaphase mainly reside on the apical side of the E8.5 neuroepithelia exposed to maternal diabetes (Fig. 2A, B). Diabetes-increased the abundance of p21, p27, and DNA damage response proteins [p-checkpoint kinase 1 (CHK1), p-CHK2, γH2AX and p53], were abrogated by *Foxo3a* deletion (Fig. S2A).

**Figure 2.**
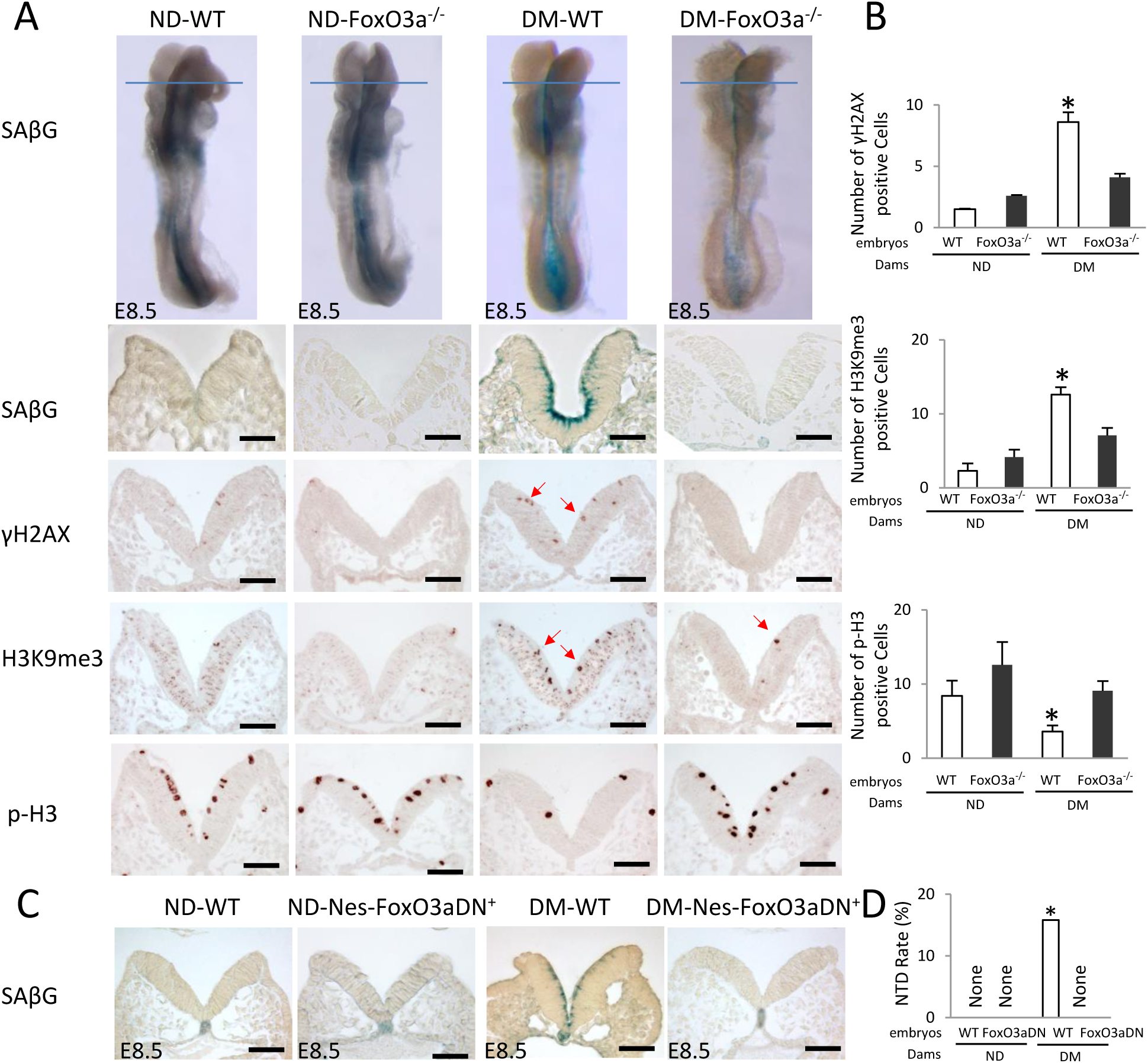
FoxO3a is essential for maternal diabetes-induced premature senescence. (A) SAβG staining and γH2AX, HeK9me3 and phosphorylated Histone 3 (p-H3) antibody staining. Blue lines indicate the levels of sectioning shown below. Quantification of antibody staining-positive cells was shown in (B). (C) Sections of SAβG staining on E8.5 WT and Nestin-Foxo3a-DN embryos. (D) NTD rates of WT and Nestin-FoxO3a-DN embryos. Scale bars = 70 μm in A, C. ND: nondiabetic dams; DM: diabetic mellitus dams; NTD: neural tube defects; WT: Wild-Type; DN: dominant negative. * indicate significant difference (*P* < 0.05) compared to other groups.

DN-FoxO3a overexpression in embryos lacking the transactivation domain from the C terminus ^32^ had a similar effect as *Foxo3a* deletion in blocking maternal diabetes-induced premature senescence (Fig. 2C). In addition, DN-FoxO3a overexpression in the neuroepithelium ablated the increase of p21, p27, p-CHK1 and p-CHK2 (Fig. S2B). Consistent with previous findings that *Foxo3a* deletion significantly reduces NTDs in diabetic pregnancy ^12^, embryos overexpressing DN-FoxO3a had a significantly lower NTD incidence compared to their WT littermates from diabetic dams (Fig. 2D, Table S1). Because DN-FoxO3a lacks transcriptional activity ^32^, the above observations implicate a transcriptional mechanism underlying FoxO3a-mediated premature senescence in diabetic embryopathy.

### Maternal diabetes-activated FoxO3a induces miR-200c expression

MicroRNAs (miRNAs) are small endogenous non-coding RNAs that suppress gene expression at the post-transcriptional level ^33^, and they play important roles in embryonic development ^34^. Recent evidence shows the critical involvement of miRNAs in mouse neurulation ^35^, NTDs in human pregnancies ^36^ and mechanisms underlying diabetic embryopathy ^11, 37^. miRNAs have been identified as novel inducers of cellular senescence ^38, 39, 40^. We hypothesized that an miRNA mediates the premature senescence in diabetic embryopathy. We focused on miR-200c because it is involved in inducing senescence in cell culture systems ^39^. Two FoxO3a binding sites were found in the promoter region of the miR-200 cluster (Fig. 3A). Constitutively active (CA) FoxO3a mimicked high glucose-induced miR-200 promoter activity (Fig. 3A), whereas DN-FoxO3a blocked the stimulatory effect of high glucose on miR-200 transcription (Fig. 3A). Either *Foxo3a* deletion or DN-FoxO3a overexpression in the developing neuroepithelium abolished the miR-200c increase in embryos of diabetic pregnancies (Fig. 3B, Fig. S3).

**Figure 3.**
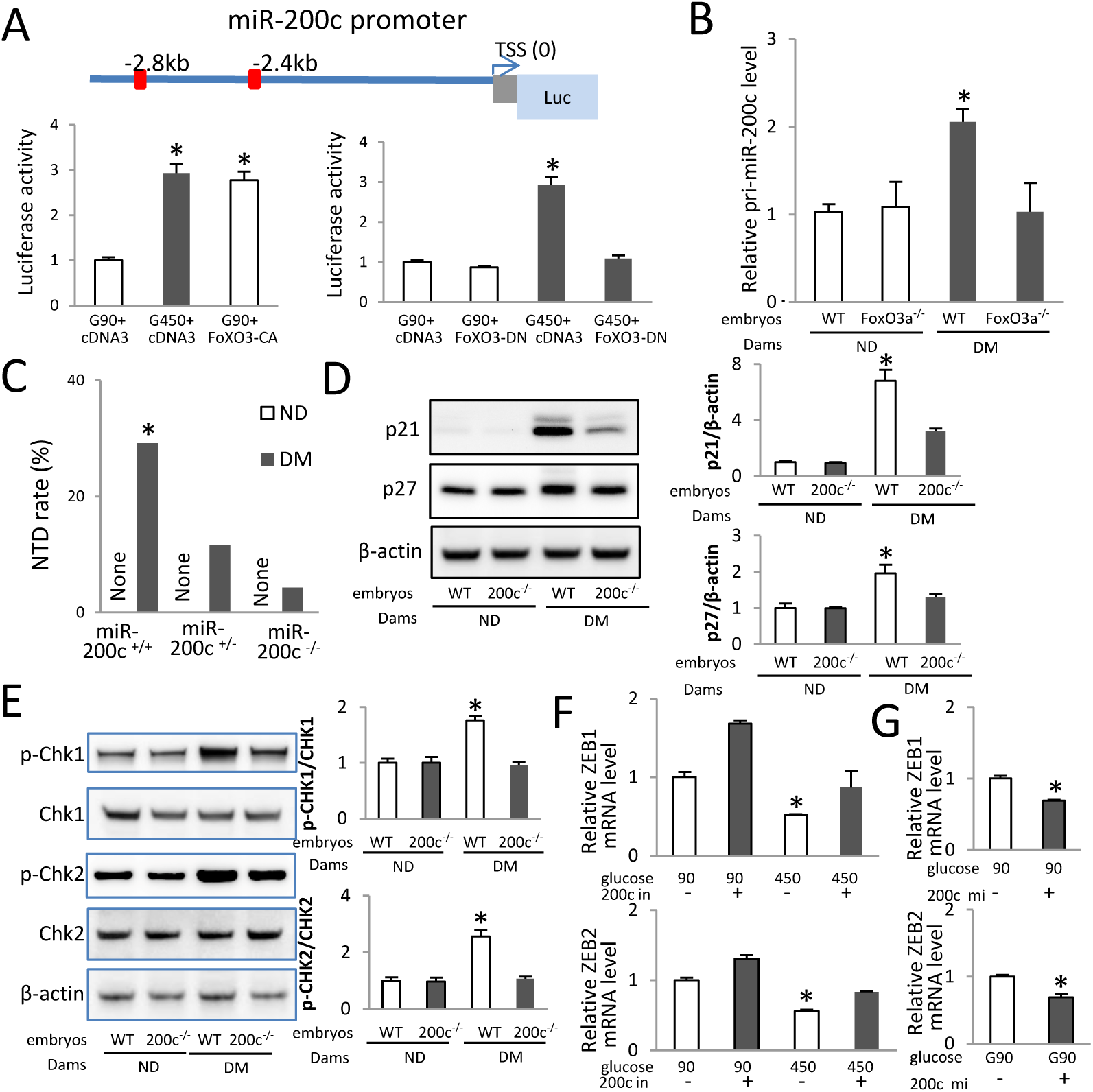
miR-200c downstream of FoxO3a is critically involved in maternal diabetes-induced premature senescence and NTDs. (A) Schematic of the miR-200 promoter-Luciferase construct. A 3.6 kb genomic fragment from the miR-200 promoter regain was cloned into the pGL4.10 vector. Red squares indicate FoxO3a binding sites. The relative Luciferase activity (Luc) after co-transfection with blank vector control, FoxO3a CA and DN expression vectors were shown. (B) pri-miR-200c abundance in WT and FoxO3a mutant embryos. (C) NTD rates of WT, miR-200c null and heterozygous embryos. Protein abundance of p21 and p27 (D); phosphorylated (p-) Chk1, Chk1, p-Chk2 and Chk2 (E) in E8.5 embryos. Bar graphs for protein abundance were quantitative data from three independent experiments. mRNA abundance of ZEB1 and ZEB2 were affected by the miR-200c inhibitor (F) and the miR-200c mimic (G) in neural stem cells (n = 3). ND: nondiabetic dams; DM: diabetic mellitus dams; NTD: neural tube defects; WT: Wild-Type. TSS: transcription starting site. * indicate significant difference (*P* < 0.05) compared to other groups.

### miR-200c mediates the effect of maternal diabetes on the induction of senescence and NTDs

To interrogate whether miR-200c plays a role in maternal diabetes-induced neuroepithelial senescence and NTD formation, heterozygous *miR-200c* KO mice were used to generate WT, *miR-200c* heterozygous and null embryos under the same maternal conditions (Table S2). *miR-200c* heterozygous embryos had an insignificant reduction of NTDs, whereas *miR-200c* null embryos achieved a significant decrease of NTD incidence compared with WT embryos from diabetic dams (Fig. 3C, Table S2). miR-141 is the closest member of miR-200c in the miR-200 family ^41^. *miR-141* was also deleted in the miR-200c KO mice (Fig. S4). To ascertain whether miR-200c is solely responsible for the NTD reduction in miR-200c null embryos, we generated *miR-200c*^*-/-*^ embryos with miR-141 transgene expression in the neuroepithelium (Table S3). Restoring miR-141 expression did not alter the inhibitory effect of *miR-200c* deletion on maternal diabetes-induced NTDs (Table S3). miR-200c overexpression in the neuroepithelium did not induce NTDs (Table S4), suggesting that other miRNAs possibly including miR-200b ^11^ co-mediate the teratogenicity of maternal diabetes with miR-200c, and miR-200c itself cannot reach the threshold leading to NTD formation. However, miR-200c overexpression neutralized the beneficial effect of *miR-200c* deletion on reducing diabetes-induced NTDs (Table S4), supporting the specificity of miR-200c deletion in NTD reduction.

To elucidate the mechanism underlying *miR-200c* deletion-reduced NTDs, indices of premature senescence in the neuroepithelium were analyzed. Maternal diabetes-increased p21, p27, p-CHK1 and p-CHK2 were significantly attenuated by miR-200c deletion (Fig. 3D, E). *miR-200c* deletion significantly decreased the signals of SAβG, γH2AX, p53 and H3K9me3, and restored the cell proliferation marker p-H3 signal in the apical side of the anterior neuroepithelium (Fig. 4A, B, C, D). These results suggest that miR-200c is responsible for upregulation of the senescence inducers-induced DNA damage signaling.

**Figure 4.**
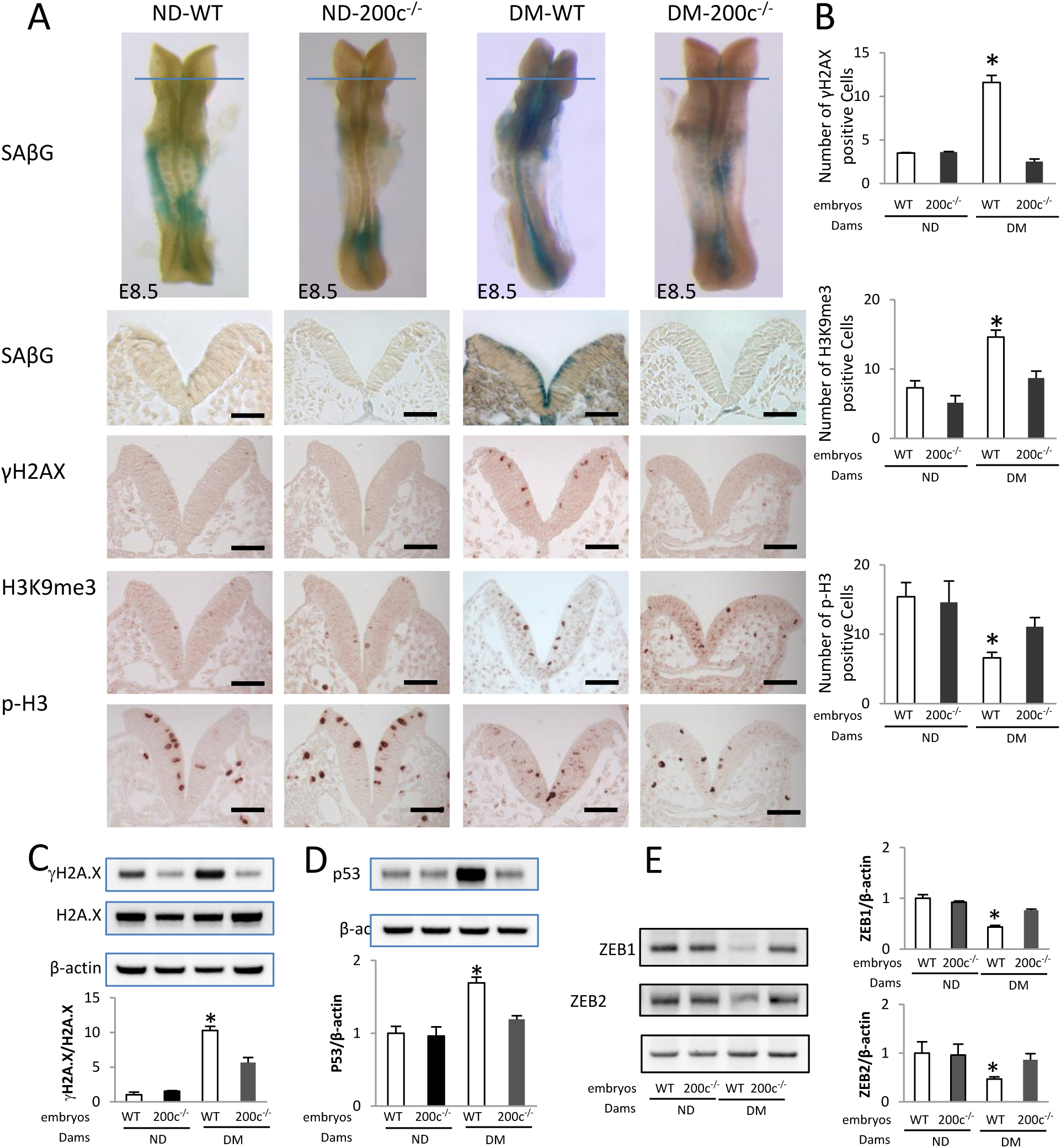
miR-200c mediates maternal diabetes induced premature senescence. (A) SAβG staining and antibody staining of gH2AX, HeK9me3 and phosphorylated Histone 3 (p-H3). Quantification of antibody staining-positive cells were shown in (B). Protein abundance of gH2A.X, H2A.X (C), p53 (D), ZEB1 and ZEB2 (E) in E8.5 embryos. Bar graphs for protein abundance were quantitative data from three independent experiments (n = 3). Scale bars = 70 µm. ND: nondiabetic dams; DM: diabetic mellitus dams; WT: Wild-Type. * indicate significant difference (*P* <0.05) compared to other groups.

### miR-200c silences transcription repressors ZEB1 and ZEB2

To search for the direct targets of miR-200c, we focused on zinc finger E-box binding homeobox 1 and 2 (ZEB1 and ZEB2), two transcription repressors for the senescence mediator, p21 ^39, 42^. An online miR target prediction algorithm (miRanda) revealed several binding sites of miR-200c to ZEB1 and ZEB2 mRNAs (Fig. S5A). To determine whether miR-200c targets ZEB1 and ZEB2 mRNA, we performed an RNA pull-down assay using biotin-labeled miR-200c. Cells transfected by biotin-labeled miR-200c had a high level of miR-200c (Fig. S5B). ZEB1 and ZEB2 mRNAs were enriched about six fold in biotin-labeled miR-200c (Fig. S5B). The miR-200c mimic simulated the effect of high glucose in repressing ZEB1 and ZEB2, whereas the miR-200c inhibitor abolished high glucose-inhibited ZEB1 and ZEB2 expression (Fig. 3F, G). Thus, miR-200c represses ZEB1 and ZEB2 expression by degrading mRNA and inhibiting translation. Consistent with the up-regulation of miR-200c, ZEB1 and ZEB2 protein abundance was significantly reduced by maternal diabetes (Fig. 4E). *miR-200c* deletion abrogated maternal diabetes-induced ZEB1 and ZEB2 down-regulation (Fig. 4E). These findings support the hypothesis that miR-200c induces premature senescence by removing the p21 and p27 promoter repressors, ZEB1 and ZEB2 ^43, 44^.

### p21 and p27 are premature senescence mediators in diabetic prenancy

To investigate whether the FoxO3a-miR-200c-ZEB1/2 pathway converges on the senescence mediators, p21 and p27, leading to failed neurulation, we used p21 KO and p27 KO mice (Table S5, 6). *p21* heterozygous embryos displayed an insignificant reduction of NTDs, whereas *p21* null embryos of diabetic dams had a significantly lower NTD incidence compared to that of WT embryos (Fig. 5A, Table S5). To validate the specificity of *p21* deletion on NTD reduction, the p21 transgene was introduced to p21 KO mice (Table S7). Restoring p21 expression in the neuroepithelium abolished the NTD reduction in *p21* null embryos of diabetic dams (Table S7). *p21* deletion reduced the SAβG, γH2AX and H3K9me3 signals, and it restored the cell proliferation marker p-H3 staining on the apical side of the anterior neuroepithelium induced by maternal diabetes (Fig. 5B, C). The induction of DNA damage markers, p-CHK1, p-CHK2, γH2AX and p53 by maternal diabetes was suppressed by *p21* deletion (Fig. 5D, E), suggesting that DNA damage response is a downstream event of p21-induced senescence. ZEB1 and ZEB2 down-regulation and miR-200c up-regulation were unaffected by *p21* deletion (Fig. 5F, G), indicating that they are upstream of p21 in the premature senescence pathway.

**Figure 5.**
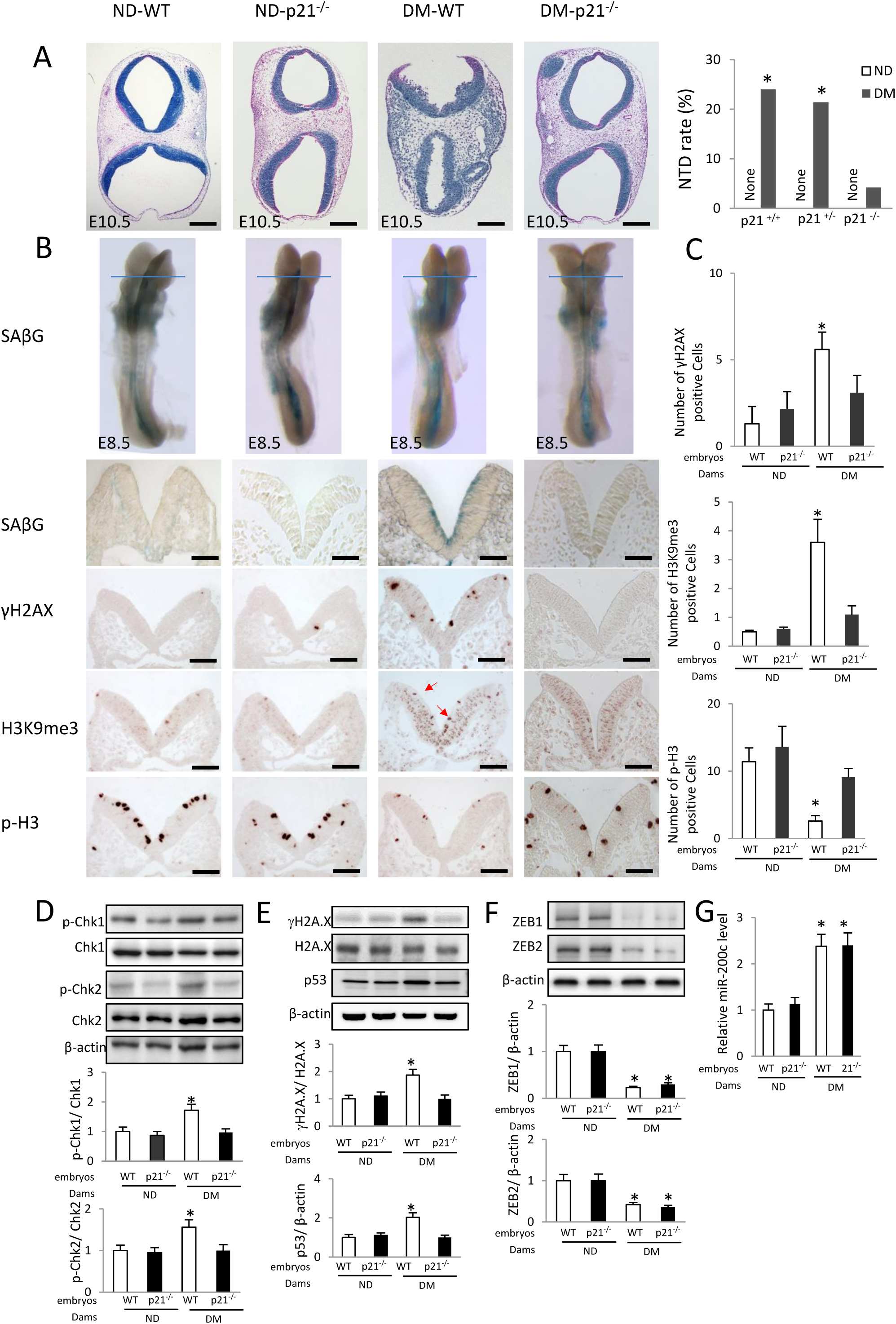
Deletion of the senescence mediators p21 abrogates premature senescence. (A) HE staining of E10.5 embryonic section to show the open neural tube in NTD embryos and NTD rates of Wild-Type (WT), p21 null and heterozygous embryos were shown. (B) SAβG staining and antibody staining of γH2AX, HeK9me3 (red arrows) and phosphorylated Histone 3 (p-H3). Quantification of antibody staining-positive cells were shown in (C). Protein abundance of phosphorylated (p-) Chk1, Chk1, p-Chk2 and Chk2 (D); γH2A.X, H2A.X and p53 (E); ZEB1 and ZEB2 (F) in E8.5 embryos. Bar graphs for protein abundance were quantitative data from three independent experiments (n = 3). (G) mRNA abundance of miR-200c in E8.5 embryos. Scale bars = 70 µm in A. ND: nondiabetic dams; DM: diabetic mellitus dams. * indicate significant difference (*P* <0.05) compared to other groups.

*p27* deletion had a similar effect as *p21* deletion, leading to a significant reduction of NTDs in diabetic pregnancies (Fig. 6A, Table S6). Heterozygous *p27* deletion slightly reduced NTD formation (Fig. 6A, Table S6). *p27* deletion effectively suppressed maternal diabetes-induced premature senescence by blocking the SAβG, γH2AX and H3K9me3 signals, and restoring cell proliferation (Fig. 6B, C). Because either *p21* or *p27* deletion significantly reduced NTDs, the additive effect of double deletion was not expected, and thus, not assessed.

**Figure 6.**
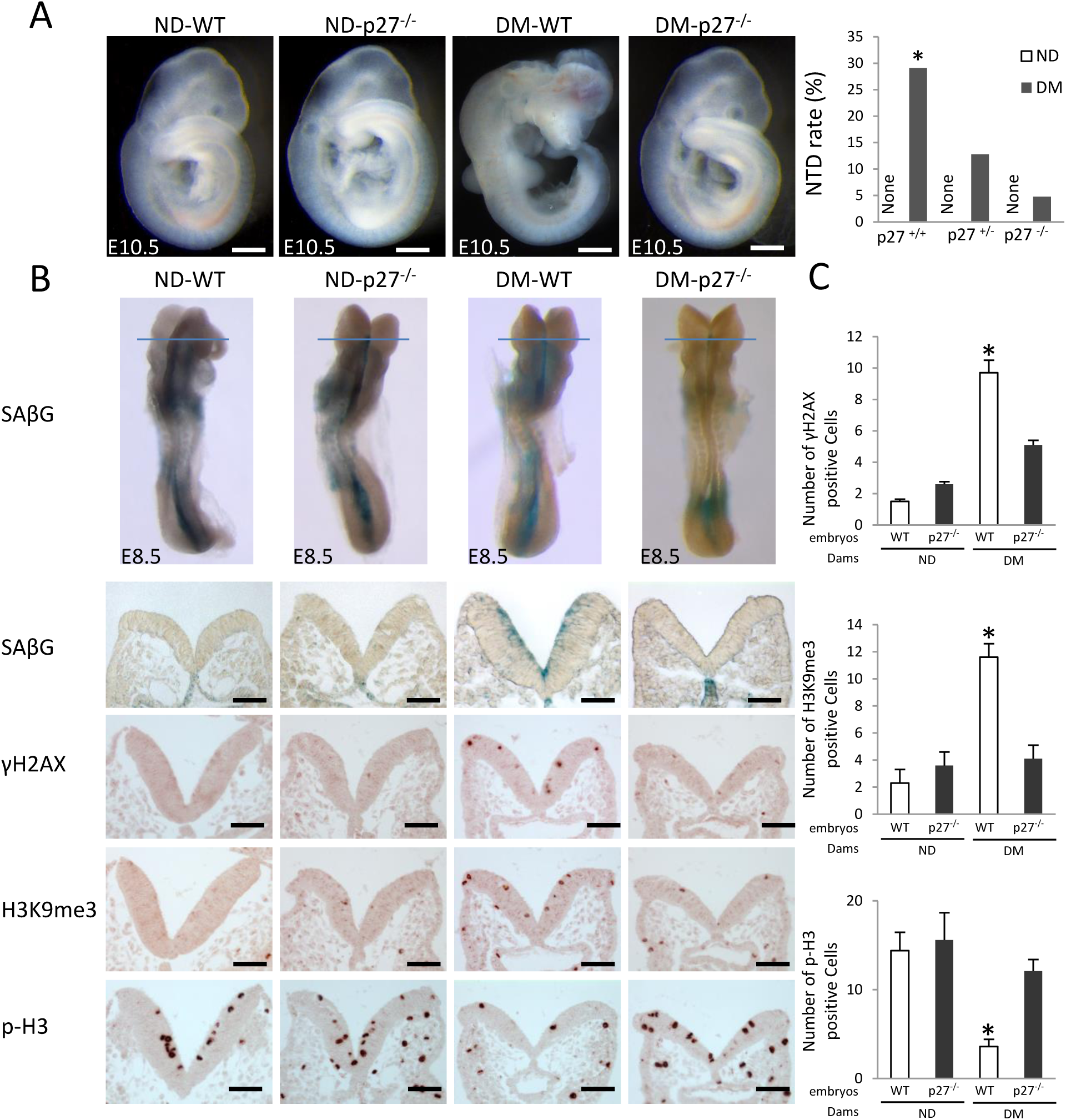
Deletion of the senescence mediator p27 abrogates premature senescence. (A) HE staining of E10.5 embryonic sections and NTD rates of Wild-Type (WT), p27 null and heterozygous embryos were shown. (B) SAβG staining and antibody staining of γH2AX, HeK9me3 and phosphorylated Histone 3 (p-H3). Quantification of antibody staining-positive cells were shown in (C). Scale bars = 70 µm in A. ND: nondiabetic dams; DM: diabetic mellitus dams. * indicate significant difference (*P* <0.05) compared to other groups.

### Double overexpression of p21 and p27 in the neuroepithelium micmics diabetes-induced NTDs

The critical involvement of both p21 and p27 in maternal diabetes-induced premature neuroepithelium senescence leading to NTD formation was exemplified by the fact that embryos with *p21* and *p27* double transgene expression in the developing neuroepithelium had a similar NTD incidence as those from diabetic pregnancies (Fig. 7A, B, Table S8). Premature senescence was manifested in the E8.5 apical side and E9.25 neural fold fusion tips of the anterior neuroepithelia of *p21* and *p27* double overexpressing embryos (Fig. 7C, D).

**Figure 7.**
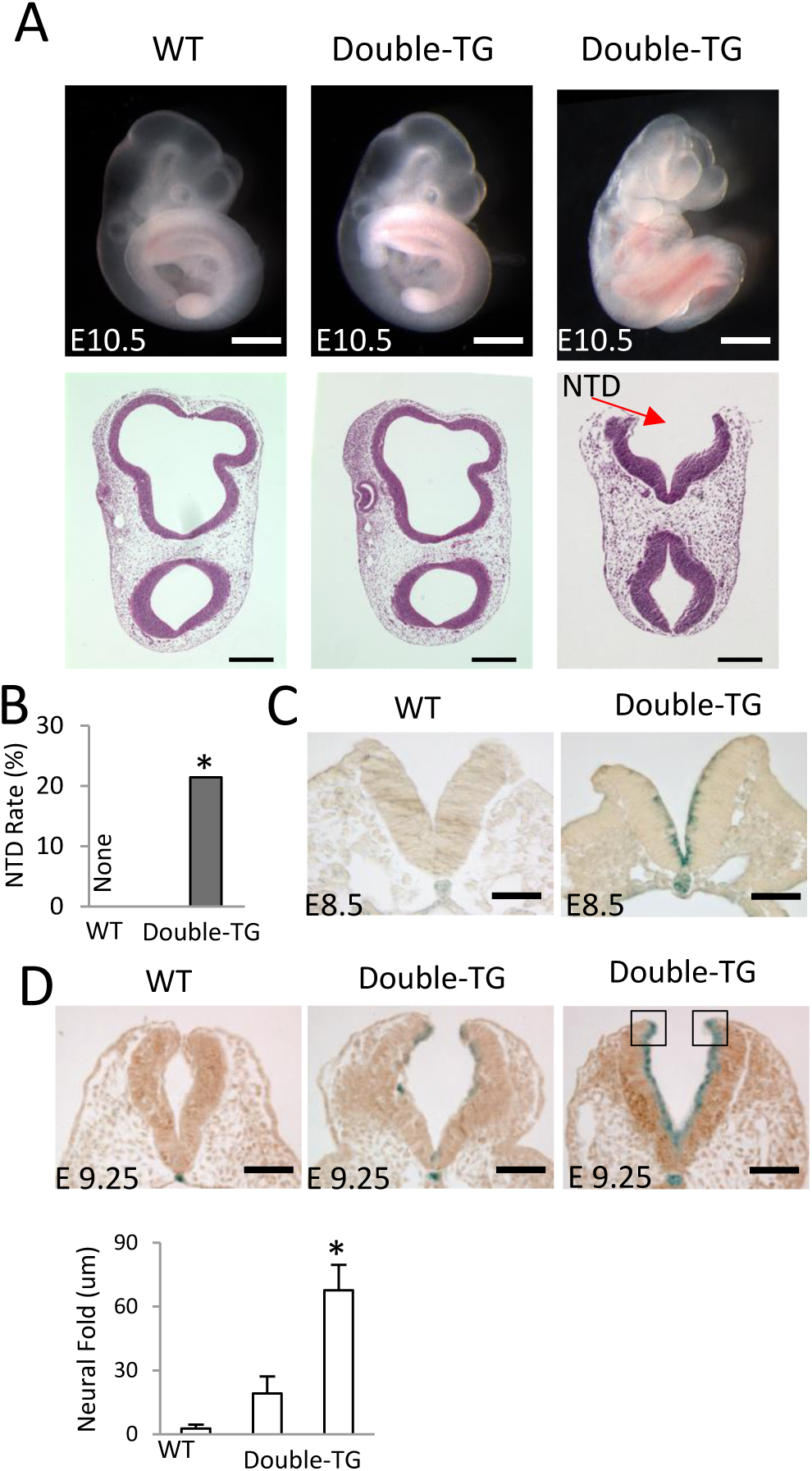
The senescence mediators, p21 and p27, mediate the teratogenicity of maternal diabetes leading to NTD formation. (A) E10.5 whole embryos and HE staining sections of WT and p21/p27 double transgenic embryos and (B) NTD rates in each group. (C) Coronal sections of E8.5 WT and p21/p27 double transgenic embryos after SAβG staining. (D) Sections of E9.25 embryos after SAβG staining, and (E) quantification of the distance between the two neural fold fusion edges (square frames). Scale bars = 70 μm in C, D; Scale bars = 200 μm in E. ND: nondiabetic dams; DM: diabetic mellitus dams; NTD: neural tube defects; WT: Wild-Type. * indicate significant difference (*P* < 0.05) compared to other groups.

### Rapamycin inhibits premature senescence and reduces NTD formation

It has been shown that the mTOR inhibitor rapamycin acts like a senomorph to suppress cellular senescence ^45^. To investigate whether rapamycin inhibits maternal diabetes-induced premature senescence in the developing neuroepithelium, pregnant WT diabetic and nondiabetic control mice received rapamycin via intraperitoneal injections at a daily dose of 2 mg/kg body weight from E5.5 to E8.5 or E10.5.

Embryos from diabetic dams treated with rapamycin achieved a significant decrease of NTD incidence when compared to embryos from diabetic dams without rapamycin treatment (Fig. 8A, B, Table S9). Rapamycin could be readily detected in the embryos of dams which received treatment (Table S9). Rapamycin treatment diminished maternal diabetes-induced SAβG signals, γH2AX and H3K9me3, and restored the cell proliferation marker p-H3 in the developing neuroepithelium (Fig. 8C, D). Rapamycin significantly reduced the number of apoptotic neuroepithelial cells induced by maternal diabetes (Fig. 8C, D). Furthermore, rapamycin blocked the increase of the senescence mediator p21; the activation of the DNA damage response proteins p-CHK1, p-CHK2, γH2AX and p53 (Fig. 8E-I); and reversed the repression of ZEB1 and ZEB2 (Fig. 8J).

**Figure 8.**
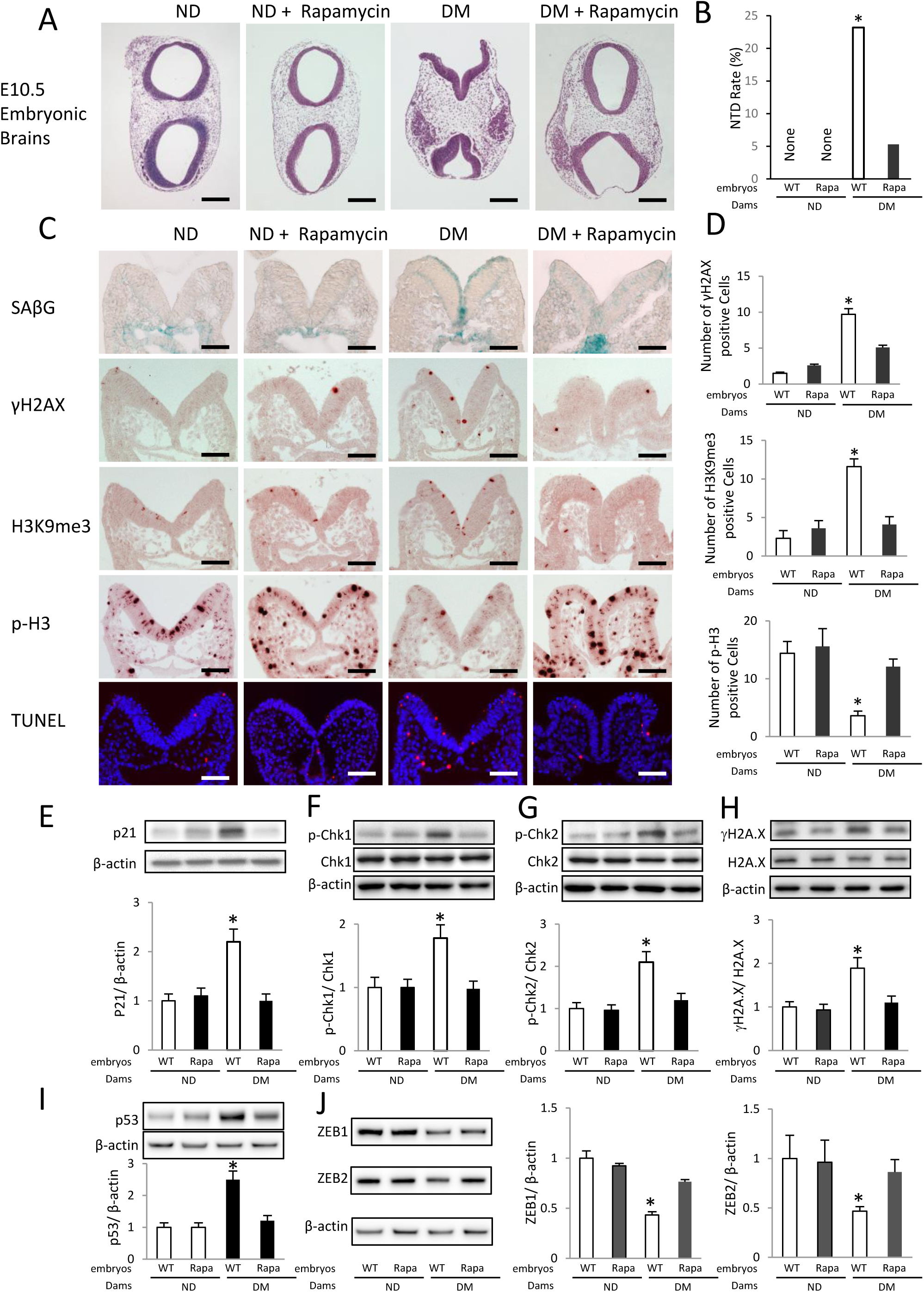
Rapamycin inhibits premature senescence and reduces NTD formation. (A) HE staining of E10.5 embryonic sections from Wild-Type dams with or without rapamycin injection. (B) NTD rates in each group. (C) SAβG staining and antibody staining of γH2AX, HeK9me3, phosphorylated Histone 3 (p-H3) and cell apoptosis detected by the TUNEL assay. Quantification of antibody staining were shown in (D). Protein abundance of p21 (E), phosphorylated (p-) Chk1 and total Chk1 (F), p-Chk2 and total Chk2 (G), γH2A.X and total H2A.X (H); p53(I) and ZEB1 and ZEB2 (J) in E8.5 embryos. Bar graphs for protein abundance were quantitative data from three independent experiments (n = 3). Scale bars = 200 µm in A and 70 µm in C. ND: nondiabetic dams; DM: diabetic mellitus dams. * indicate significant difference (*P* < 0.05) compared to other groups.

## Discussion

Senescence as a new mechanism for human adulthood diseases has just been recognized. Drugs that target senescent cells, known as senotherapeutics, work by inhibiting the initiation of senescence (senomorphic drugs) or by removing senescent cells (senolytic drugs) have just been tested in small-scale clinical trials ^46, 47^. In the current study, we revealed that premature senescence in the developing neuroelithium contributes to the induction of NTDs caused by maternal diabetes in pregnancy. This form of premature senescence is initiated by transcriptional and epigenetic alterations of neurogenesis via the FoxO3a-miR-200c-ZEB1/2-p21/p27 pathway. Based on these findings, inhibition of premature senescence could be a potential intervention for diabetic embryopathy. It has been shown that rapamycin can inhibit cell entry into senescence ^45^. We showed that rapamycin inhibited diabetes-induced premature senescence in the developing neuroepithelium and reduced the number of apoptotic neuroephtelial cells, suggesting that rapamycin acts as a senomorph that inhibits the iniation of senescence. Rapamycin is an FDA-approved drug in preventing the immune-rejection of transplanted organs with a good safety profile ^48^ and thus, it may be repurposed to treate diabetes-induced embryonic malformations as a senomorph.

Successful neurulation requires a critical cell mass in the dorsal lateral hinge points (DLHP), which induce the inward bending and eventual fusion of the two dorsal neural fold tips to creat the neural tube ^28, 29^. Neuroepithelial cell proliferation in the apical side of the neuroepithelium contributes to the critical cell mass in the DLHPs through cell migration in a ventral-to-dorsal direction ^28, 29^. Here, we observed that maternal diabetes targets apical neuroepithelial cells undergoing M-phase in mitosis and induces these cells to undergo premature senescence, thereby reducing the cell mass in the DLHPs required for neural tube closure. In addition, we found that senescent cells may adversely impact other neuroepithelial cells through the SASP paracrine mechanism.

The degree of senescence in the apical neuroepithelium ranges from modest to severe. Modest senescence may not have any NTD manifestation and may result in other developmental defects such as microcephaly ^13^, autism and other development disabilities ^49^. Severe senescence results in NTD formation. In addition to apical neuroepithelial cell senescence, the occurrence of senescence in the cells comprsing the tips of the two dorsal neural folds during the late phase of neurulation may also be required for NTD formation. Embryos with senescent cells in the dorsal neural fold tip fail to properly bend the DLHPs and display wider gaps between the two neural fold tips than embryos without senescent cells in this area.

The current study provides genetic evidence supporting a causal role of the FoxO3a-miR-200c-ZEB1/2-p21/p27 pathway in maternal diabetes-induced apical neuroepithelial cell premature senescence and NTD formation. Deletion of the *Foxo3a* gene, the *miR-200c* gene, the *p21* gene or the *p27* gene abrogated senescence and ameliorated NTDs in embryos of diabetic dams. Importantly, *p21* and *p27* transgenic overexpression induced apical neuroepithelial cell premature senescence and NTDs, similar to maternal diabetes. These findings collectively support our hypothesis that premature senescence in the apical neuroepithelium causes NTDs in diabetic pregnancies.

FoxO transcription factors regulate a variety of cellular functions including cell proliferation, apoptosis, aging and oxidative stress. In adult cells, FoxO family members differentially regulate senescence ^50, 51^. As a tumor suppressor, FoxO4 induces p21-dependent senescence in cancer cells ^52^. In contrast, FoxO3a suppresses senescence in aging cells ^51^. Therefore, FoxO factors regulate senescence in a cell-type and cellular context-dependent manner ^50^. Recent evidence support the hypothesis that FoxO factors are senescence inducers during embryonic development ^19^. During normal embryogenesis, FoxO1 is the main stimulator of developmental senescence ^19^. In diabetic embryopathy, previous studies have demonstrated that maternal diabetes activates FoxO3a, but not FoxO1 or FoxO4 ^12^. Both *Foxo3a* deletion and transgenic overexpression of DN-FoxO3a blocked maternal diabetes-induced apical neuroepithelial senescence and NTD formation, strongly supporting FoxO3a as the senescence inducer in diabetic embryopathy.

FoxO factors could directly induce senescence through their transcription activity by inducing cell cycle inhibitors ^52^. However, the present study reveals that FoxO3a triggers neuroepithelial cell senescence through up-regulation of miR-200c. Indeed, recent evidence implies that FoxO factors differentially regulate miRNA expression ^53, 54^. We observed that deleting Foxo3a or overexpressing DN-FoxO3a in the neuroepithelium abolished the induction of miR-200c, supporting the concept that FoxO3a transcriptionally up-regulates miR-200c. miR-200c belongs to the miR-200 family, which consists of five members including miR-141 and miR-200b. Both miR-200c and miR-141 can induce cellular senescence *in vitro* ^38, 39^. miR-141 overexpression does not affect the preventive effect of miR-200c deletion on diabetic embryopathy. Therefore, only miR-200c appears to be involved in maternal diabetes-induced premature senescence in the developing neuroepithelium. This differential effect of miR-141 and miR-200c may be explained by their different seed sequences. Indeed, miR-141 induces senescence by repressing the polycomb gene BMI1, leading to the increase of the cell cycle inhibitor, p16 ^38^. In contrast, miR-200c silences ZEB1 and ZEB2, which remove the repression of p21 and p27, leading to premature senescence in the developing neuroepithelium.

p21 and p27 cooperatively induce senescence in the developing neuroepithelium. Individual deletion of either the *p21* gene or the *p27* gene abrogates premature senescence in diabetic embryopathy. p21 not only mediates developmental senescence but also cellular senescence induced by a variety of stresses ^55^. In agreement with the present finding that p21 gene deletion blocks the induction of premature senescence in diabetic embryopathy, ablation of p21 prevents both developmental senescence *in vivo* and stress-induced senescence *in vitro* ^19, 56^. p27 is the closest cell cycle inhibitor to p21 and they belong to the same family of cell cycle inhibitors ^57^. Studies have also demonstrated that p27 contributes to senescent phenotypes ^58^. Furthermore, p27 directly induces senescence in adult cells ^59, 60^. The evidence collectively support the importance of both p21 and p27 in the induction of premature senescence in diabetic embryopathy. Both the p21 KO mouse model and the p27 KO model are ideal for the present study because deletion of either the *p21* gene or the *p27* gene does not affect neurulation and overall embryonic development ^61, 62^. Although p27 KO female mice have reproductive problems ^62^, the study has circumvented these problems by effectively using the heterozygous mating scheme. The most striking finding from this study is that overexpressing both p21 and p27 mimicks maternal diabetes in inducing senescence in the apical neuroepithelium, leading to NTD formation.

Rapamycin is an inhibitor of mTOR, whose activation leads to cell senescence ^63^. A previous study suggested that mTOR inhibition suppresses miR-200c expression and increases ZEB1/2 expression during epithelial-to-mesenchymal transition ^64^. Combined with the present results, mTOR and FoxO3a may cooperatively upregulate miR-200c leading to premature senescence in diabetic embryopathy. Furthermore, the mTOR effect may be mediated by its downstream effector, p70S6K1, whose activation critically participates in the induction of diabetic embryopathy ^65^.

In summary, the present study is the first one to reveal a critical role for premature senescence in the apical neuroepithelium in diabetic embryopathy. The transcription factor, FoxO3a, activated by oxidative stress ^12^, triggers a senescence pathway that involves miR-200c, ZEB1/2, p21 and p27. The apical proliferative metaphase neuroepithelial cells are primary targets of premature senescence in diabetic pregnancy, and senescence in these cells adversely impact the neurulation process by inhibiting neural fold bending and tip fusion. Thus, premature senescence is a new mechanism underlying failed neural tube closure in diabetic embryopathy. Inhibiting senescence using the the senomorph-like agent rapamycin is an effective method to prevent diabetic embryopathy.

## Methods

### Mice

All procedures for animal use were approved by the Institutional Animal Care and Use Committee of University of Maryland School of Medicine. Wild-type (WT) C57BL/6J mice, p21 KO, p27 KO and FoxO3a KO mice were purchased from the Jackson Laboratory (Bar Harbor, ME). The miR-200c/141 knockout (KO) mouse in a C57BL/6J background was made in the Keio University School of Medicine (Fig. S4) with a similar strategy for the miR-200b/429 knockout mice ^66^. The male heterozygous deletion p21^+/-^ or p27^+/-^ mice were mated with nondiabetic or diabetic female p21^+/-^ or p27^+/-^ mice to generate Wild-Type (WT), p21^+/-^ or p27^+/-^, p21^-/-^ or p27^-/-^ embryos. The male FoxO3a^+/-^ mice were mated with nondiabetic or diabetic female FoxO3a^+/-^ mice to generate WT, FoxO3a^+/-^, and FoxO3a^-/-^ embryos. The male miR-200c/141^+/-^ mice were mated with nondiabetic or diabetic female miR-200c/141^+/-^ mice to generate wild type (WT), miR-200c/141^+/-^, and miR-200c/141^-/-^ embryos.

### Generation of DN-FoxO3a, miR-200c, miR-141, p21 and p27 transgenic mice

The dominant negative FoxO3a (DN-FoxO3a), miR-200c, miR-141, p21 and p27 transgenic (Tg) mice were generated by pronuclear injections using the standard procedures in the Genome Modification Facility, Harvard University. The coding region of DN-FoxO3a was cloned from the Addgene plasmid #1797 ^67^, a gift from Michael Greenberg. The full-length mouse complementary DNAs (cDNA) encoding pre-miR-200c, pre-miR-141, p21 and p27 with a separated expressed GFP were subcloned to the downstream of rat nestin promoter and followed by a 3’ nestin genomic sequence as previously described ^68^. The transgenic constructs were linearized to remove vector sequence, injected into fertilized oocytes from C57BL/6 female mice, and implanted into pseudopregnant mice. 3-4 founders of each Tg line were generated and founders with 2-4 fold transgene expression were selected for experiments.

### Model of diabetic embryopathy

We ^7, 8, 11, 12, 69, 70, 71^ and others ^72, 73, 74^ have used a rodent model of Streptozotocin (STZ)-induced diabetes in research of diabetic embryopathy. Briefly, ten-week old WT or knockout female mice were intravenously injected daily with 75 mg/kg Streptozotocin (STZ) for two days to induce diabetes. STZ from Sigma (St. Louis, MO) was dissolved in sterile 0.1 M citrate buffer (pH4.5). We used the U-100 Insulin Syringe (Becton Dickinson) with 28G1/2 needles for injections. Approximately 140 µl of STZ solution was injected per mouse. Diabetes was defined as a 12-hour fasting blood glucose level of ≥ 16.7 mM. Male and female mice were paired at 3:00 P.M., and day 0.5 (E0.5) of pregnancy was established at noon of the day when a vaginal plug was present. Embryos were harvested for subsequence analysis on different developing stage. Embryos were harvested at 2:00 PM at E8.5 for biochemical and molecular analyses. At E10.5, embryos were examined under a Leica MZ16F stereo microscope (Bannockburn, IL) for NTD examination. Images of embryos were captured by a DFC420 5 MPix digital camera with software (Leica, Bannockburn, IL) and processed with Adobe Photoshop CS2. Normal embryos were classified as possessing completely closed neural tube and no evidence of other malformations. Malformed embryos were classified as showing evidence of failed closure of the anterior neural tubes resulting in exencephaly, a lethal type of NTDs with the absence of a major portion of the brain, skull, and scalp. Because our model induces NTD at E10.5, we did not examine other major structural malformations such as cardiovascular defects, which do not occur until later embryonic stages (E13.5).

### Rapamycin treatment to pregnant mice

Rapamycin was dissolved in DMSO to a concentration of 25mg/ml and stored at -20 °C. This stock solution was diluted by 50 X to a concentration of 0.5mg/ml with vehicle containing 5% Tween-80 and 30% PEG400. Pregnant Wild-Type diabetic and nondiabetic mice received intraperitoneal injections of rapamycin at a dose of 0.1 ml/g body weight (2 mg/kg body weight) based on a previous study ^75^. Pregnant mice in the control groups (diabetic and nondiabetic) received the same volume of the vehicle solution. Rapamycin injections was given daily from E5.5 to E8.5 for biochemical and molecular analyses and to E10.5 for morphological examination.

### Whole mount senescence staining

For whole mount embryo staining, embryos were fixed overnight in 0.5% gluteraldehyde prepared with PBS (pH = 7.4). Embryos were washed with PBS/MgCl_2_ (pH = 5.5, 1 mM MgCl_2_) for 2 × 15 min at 4°C on a rocker, and then stained with the X-gal staining solution containing 0.25 ml of X-gal Stock (Roch #745740) in N,N-dimethylformamide (sigma #D-4254)), 0.25 ml of 0.2 M K_3_Fe(CN)_6_, 0.25 ml of 0.2 M K_4_Fe(CN)_6_·3H_2_O and 9.25 ml of PBS/MgCl_2_ for 4 hours. For embryo sectioning after senescence staining, embryos were embedded in OCT compound and sectioned (10 μm) and mounted on superfrost slides for microscopy analysis.

### Neuroepithelial cell isolation and flow cytometry analysis of senescent cells

E8.5 embryos were dissected out of uterus and then yolk sacs were removed from the embryos. The whole embryos were digested by Dispase II (3 mg/ml in PBS pH7.4) for 15 min at room temperature in order to remove non-neural tissues ^76, 77^. After the enzymatic digestion, whole neuroepithelia were washed with PBS for three times, cut into pieces and digested with 0.25% trypsin-EDTA (Thermo Scientific) for 5 min at 37°C with shaking. The cell suspensions were centrifuged at 500 g for 5min and resuspended in DMEM medium (Thermo Scientific) with 10% fetal bovine serum (FBS) (Thermo Scientific) following by filtration through a 70 μm cell strainer. The isolated cells were seeded into 6-well plate.

The live cell senescence analysis was performed using the Cellular Senescence Live Cell Analysis Assay Kit (Enzo Life Sciences). Briefly, after adhesion, the isolated neuroepithelial cells were treated with 2 ml of 1× cell pretreatment solution for 2 hours at 37°C. Then 10 μl of 200× senescence-associated □-galactosidase (SA□G)substrate solution was directly added into 1× cell pretreatment solution, gently mixed and incubated at 37°C for 4 hours. After senescence staining, the cells were washed with PBS for three times. The stained cells were fixed and permeabilized by the Fixation and Permeabilization Solution (BD Biosciences) following by immunostaining using a p21antibody (Table S10.) and a fluorescence-coupled secondary antibody. Finally, cells were washed and analyzed by a flow cytometer.

### Immunostaining and Hematoxylin-Eosin (HE) staining

Various stage embryos were fixed in methacarn (60% methanol, 30% chloroform, and 10% glacial acetic acid), embedded in paraffin, and cut into 5 μm sections. After deparaffinization and rehydration, all specimens were subjected to immunofluorescent staining for p21, p27, and immunohistochemical staining for γH2AX, H3K9me3, phosphorylated Histone 3 (p-H3) and F4/80 using respective antibodies (Table S10.) or HE staining. All sections were photographed.

### Cell culture and miR-200c mimic and inhibitor transfection

C17.2 mouse neural stem cells, originally obtained from ECACC (European Collection of Cell Culture), were maintained in DMEM (5 mM glucose) supplemented with 10% FBS, 100 U/ml penicillin and 100 µg/ml streptomycin at 37°C in a humidified atmosphere of 5% CO_2_. Although the C17.2 cells are newborn mouse cerebellar progenitor cells transformed with retroviral v-myc ^78^, these cells mimic the *in vivo* neuroepithelial cells in responding to high glucose ^71^. Lipofectamine RNAiMAX (Invitrogen) was used according to the manufacturer’s protocol for transfection of the miR-200c mimic and inhibitor into the cells under 1% FBS culture condition. The mirVana® miR mimic and the miR inhibitor of miR-200c were purchased from Ambion.

### MiR-200 promoter activity analysis

The binding sites of transcription factor FoxO3a which potentially regulate miR200 promoter activity were predicted using the online prediction tools (http://jaspar.genereg.net). We identified two FoxO3a binding sites in the miR-200 promoter region. The miR-200 promoter was subcloned and inserted into the pGL4.10 Luciferase Reporter Vectors (Promega). Co-transfection of FoxO3a constituted active (CA-FoxO3a-CA) or dominant negative (DN-FoxO3a) vector and the pGL4.1-miR200 promoter vector into the mouse neural stem cell line C17.2 was performed using Lipofectamine 2000 (Invitrogen) and the cells were cultured for 48 hours with or without treatment of high glucose (25 mM). CA-FoxO3a vector (addgene plasmid #1788) and DN-FoxO3a vector (addgene plasmid #1797) were gifts from Michael Greenberg ^67^. Cells were harvested and lysed using the lysis buffer of the Dual luciferase Assay system (Promega). Firefly luciferase activities were normalized to the internal control *Renilla* luciferase.

### Immunoblotting

Equal amounts of protein from embryos were resolved by the SDS-PAGE gel electrophoresis and transferred onto Immobilon-P membranes (Millipore). Membranes were incubated in 5% nonfat milk for 45 minutes at room temperature and then incubated for 18 hours at 4°C with the following primary antibodies diluted into 5% nonfat milk: p21, p27, phospho-Chk1, Chk1, phospho-Chk2, Chk2, ZEB1, ZEB2, p53, γH2AX (Table S10). Membranes were then exposed to goat anti-rabbit or anti-mouse secondary antibodies. To confirm that equivalent amounts of protein were loaded among samples, membranes were stripped and probed with a mouse antibody against β-actin (Table S10). Signals were detected using the SuperSignal West Femto Maximum Sensitivity Substrate kit (Thermo Scientific). All experiments were repeated three times with the use of independently prepared tissue lysates.

### RT-qPCR

Total RNA was isolated from tissues and cells using the TRIzol reagent (Ambion) and reverse transcribed using the QuantiTect Reverse Trancription Kit (Qiagen). Reverse transcription for miRNA was performed using the qScript microRNA cDNA Synthesis Kit (Quanta Biosciences). RT-qPCR of mRNA, pri-miRNA and small nuclear RNA U6 was performed using the Maxima SYBR Green/ROX qPCR Master Mix assay (Thermo Scientific). The primers for RT-qPCR are listed in Table S11. RT-qPCR and subsequent calculations were performed by a StepOnePlus™ Real-Time PCR System (Applied Biosystem).

### miRNA pulldown

The biotin-labeled miR-200c and its control (*C. elegans* miR-67) were custom made by GE Dharmacon. The biotin-labeled miR-200c and biotin-labeled negative control were transfected using Lipofectamine 2000 (Invitrogen) into C17.2 cells for 48 h, and then cells were lysed in a lysis buffer (20 mM Tris-HCL (pH 7.5), 100 mM KCL, 5 mM MgCl_2_, 0.3% IGEPAL CA-630). Cell lysates were mixed with streptavidin-coupled Dynabeads (Invitrogen) and incubated at 4°C on rotator overnight. After the beads were washed thoroughly, the bead-bound RNA was isolated and subjected to RT followed by Real-time PCR analysis.

## Statistical analyses

All experiments were completely randomized designed and repeated in triplicate. Data are presented as means ± standard errors (SE). Student’s t test was used for two group comparisons. One-way ANOVA was performed for more than two group comparisons using the SigmaStat 3.5 software. In ANOVA analysis, a *Tukey* test was used to estimate the significance. Significant differences between groups in NTD incidence expressed by number of embryos were analyzed by Chi-square test or Fisher’s exact test using SigmaStat 3.5 software. Statistical significance was indicated when *P* < 0.05.

## Author Contributions

C. Xu researched and organized the data. Peixin Yang conceived the project, designed the experiments, and wrote the manuscript. H. Hasuwa created the miR-200C KO mice and reviewed the manuscript. E. A. Reece, C. Harman, S. Kaushal and W.B. Shen analyzed and reviewed the manuscript.

## Acknowledgments

This work was supported by the NIH grants NIH R01DK083243, R01DK101972, R01HL131737, R01HL134368, R01HL139060 and R01DK103024. We appreciate the Genome Modification Facility, Harvard University, in generating the FoxO3a-DN transgenic (Tg), miR-141 Tg, miR-200c Tg, p21 Tg, and the double p21/p27 Tg mice. We thank Dr. Julie A. Rosen at the University of Maryland School of Medicine for critical review and editing assistance, Xi Chen at the University of Maryland School of medicine for assisting on the cell sorting analysis and Daoying Dong at the University of Maryland School of medicine for assisting the miR-200c study.

